# Variability of Phenylalanine side chain conformations facilitates promiscuity of Fatty acid binding in Cockroach milk proteins

**DOI:** 10.1101/2022.12.21.521413

**Authors:** Partha Radhakrishnan Santhakumari, KanagaVijayan Dhanabalan, Saniya Virani, Amber S. Hopf-Jannasch, Joshua B. Benoit, Gaurav Chopra, Ramaswamy Subramanian

## Abstract

The pacific beetle cockroach, *Diploptera punctata*, is a viviparous cockroach that produces a milk-like substance to support the growing embryo with a brood sac. The structure of the *in vivo* grown crystals present in the gut of the embryo showed that the milk-derived crystals are heterogenous and are made of three proteins (called Lili-Mips). Multiple fatty acids could be modeled into the active site, and we hypothesized that each of the three isoforms of the protein bound to a different fatty acid. We previously reported that the recombinantly expressed Lili-Mip2 has a structure similar to the structure of the protein determined from *in vivo* crystals, and this single isoform also binds to several fatty acids. In this study, we aimed to probe the specificity and affinity of fatty acid binding and test the stability of different isoforms. We show that all the isoforms can bind to different fatty acids with very similar affinities, and the local abundance of a fatty acid determined bound fatty acid ratios. Lili-Mips’ thermostability is pH dependent, where stability is highest at acidic pH and declines as the pH increases to physiological levels near 7.0. The measurement of the pH in the gut lumen and the gut cells suggests that the pH in the gut is acidic and the pH inside the gut cells is closer to neutral pH. We propose that the protein has evolved to be highly stable in the acidic gut lumen and, when absorbed inside the gut cells, becomes less stable to enable the breakdown of the glycosylated lipo-protein complex to provide essential metabolites for survival and development of the embryo. The different orientations of Phe-98 and Phe-100 control the binding pocket volume and allow the binding of different chain-length fatty acids to bind with similar affinities.

## 1 Introduction

The early ancestors of cockroaches lived as early as 300 million years ago [1]. Their fossils date back to 120 million years ago [2]. The robustness of this species is exemplified by its resistance to insecticides and its ability to adapt to its environment [3]. Cockroaches have evolved a reproductive nature that can be oviparous, ovo-viviparous, or viviparous [4,5]. *Diploptera punctata* (pacific beetle cockroach), the only known viviparous cockroach, gives birth and nourishes its young ones [5]. The mother secretes a concoction rich in proteins, carbohydrates, and lipids called cockroach milk while the embryos develop within the brood sac. The principal component of this milk is lipocalin-like milk proteins or Lili-Mip [6–8]. There are 25 cDNA sequences that code for Lili-Mips, expressing at least 22 unique but similar proteins [7,8]. In the embryo midgut, post-ingestion, Lili-Mip crystallizes [6]. The structure of the crystals showed that at least three of the Lili-Mips are present in a single crystal [8]. These sequences, called Lili-Mip1, 2, and 3, are highly expressed during late lactation [9]. These proteins are observed to form *in vivo* crystals despite having heterogeneity in sequence, glycosylation, and the bound ligand [6–8].

Lili-Mip has a lipocalin fold [8]. The secondary structure comprises an alpha helix and eight antiparallel beta sheets. Lili-Mip is small (18.8 kDa), monomeric and glycosylated. The calyx, or the ligand binding site, is encased by the beta sheets and resembles a barrel. Its binding pocket can accommodate different fatty acids. Crystal structure and mass spectrometric studies suggested that the bound fatty acid is either palmitoleic, oleic, or linoleic [8,10]. Ligand promiscuity in lipocalins is hypothesized to be a result of the flexibility of the loops connecting the beta sheets [11]. Lipocalins can bind to various molecules, including fatty acids, phospholipids, glycolipids, and stearic acid [12–14].

The binding and delivery of hydrophobic ligands to different places are critical parts of lipocalin function. Apolipoprotein D is a classic example of a lipid transporter [15]. The ligand binding and release mechanism of different lipocalins has been investigated. Human retinoic acid binding protein and tear lipocalin are known to release ligands at an acidic pH [16,17].

Lipocalins show nano-to-micromolar binding affinities to various hydrophobic ligands [12,18]. Many lipocalins bind to multiple ligands [18]. This is, in turn, determined by the ligand binding site and the four loops at the entrance of the calyx [13]. The large barrel that houses the ligand makes lipocalins amenable for protein engineering [8,13]. These four loops that connect antiparallel beta sheets can be designed to target the lipocalin to bind to any molecule of choice – haptens, fatty acids, peptides, and so on. These modified lipocalins are called Anticalins [19]. In tear lipocalin and Lili-Mip 2, Zinc is involved in crystal packing [10,13]. Zinc binds at the entrance of the calyx and is responsible for the preferential binding of Lili-Mip 2 to palmitoleic acid rather than oleic acid [10].

The remarkable structural similarity of lipocalins warrants the conservation of some unique sequential features. For instance, the overall RMS deviation values of Lili-Mip (PDB: 4NYQ) with respect to Human tear Lipocalin(PDB: 1XKI) is 4.5 Å for 103 C-alpha atoms [8]. The disulfide bonds and tryptophan at the base of the β-barrel are the most conserved [8,13,20,21]. The tryptophan at the N terminus (W20 in Lili-Mip) covers the base of the 10 Å wide and 15 Å deep ligand binding calyx [8,10]. Tryptophan at this position is known to be important for the structure and preventing oxidation of retinol in β-lactoglobulin. It stabilizes the protein and plays a role in binding [12,21]. The presence of this single tryptophan makes Lili-Mip, like many other members of its family, amenable to binding studies through the measurement of intrinsic tryptophan fluorescence [12,18].

Structures of Lili-Mip and Lili-Mip2 provide a molecular basis for fatty acid binding [8,10]. The Glutamate(E38) at position 38 forms a kink in the ligand binding site in Lili-Mip2 (Figure 1). It is held in position by histidine (H115) and tyrosine (Y40). One could hypothesize that it favors the binding of Lili-Mip2 to omega-6 unsaturated fatty acids [10]. In tear lipocalin, charged residues Glutamate 34, Histidine 84, and Lysine 114 are thought to be playing a role in ligand interaction [13]. In Lili-Mip, the carboxylic acid moiety of the fatty acid is exposed to the surface, whereas the acyl chain is buried [8,10]. This has been observed for another lipocalin-like retinol-binding protein, RBP4 [22], that binds laurate, palmitate, oleate, and linoleate. Glutamate 38 and Phenylalanine at 98 and 100 are conserved between these structures [8,10,22,23]. However, the carboxylate group faces inwards and interacts with tyrosine and arginine in the proteins of the fatty acid binding proteins (FABPs) family [24].

**Figure 1.**
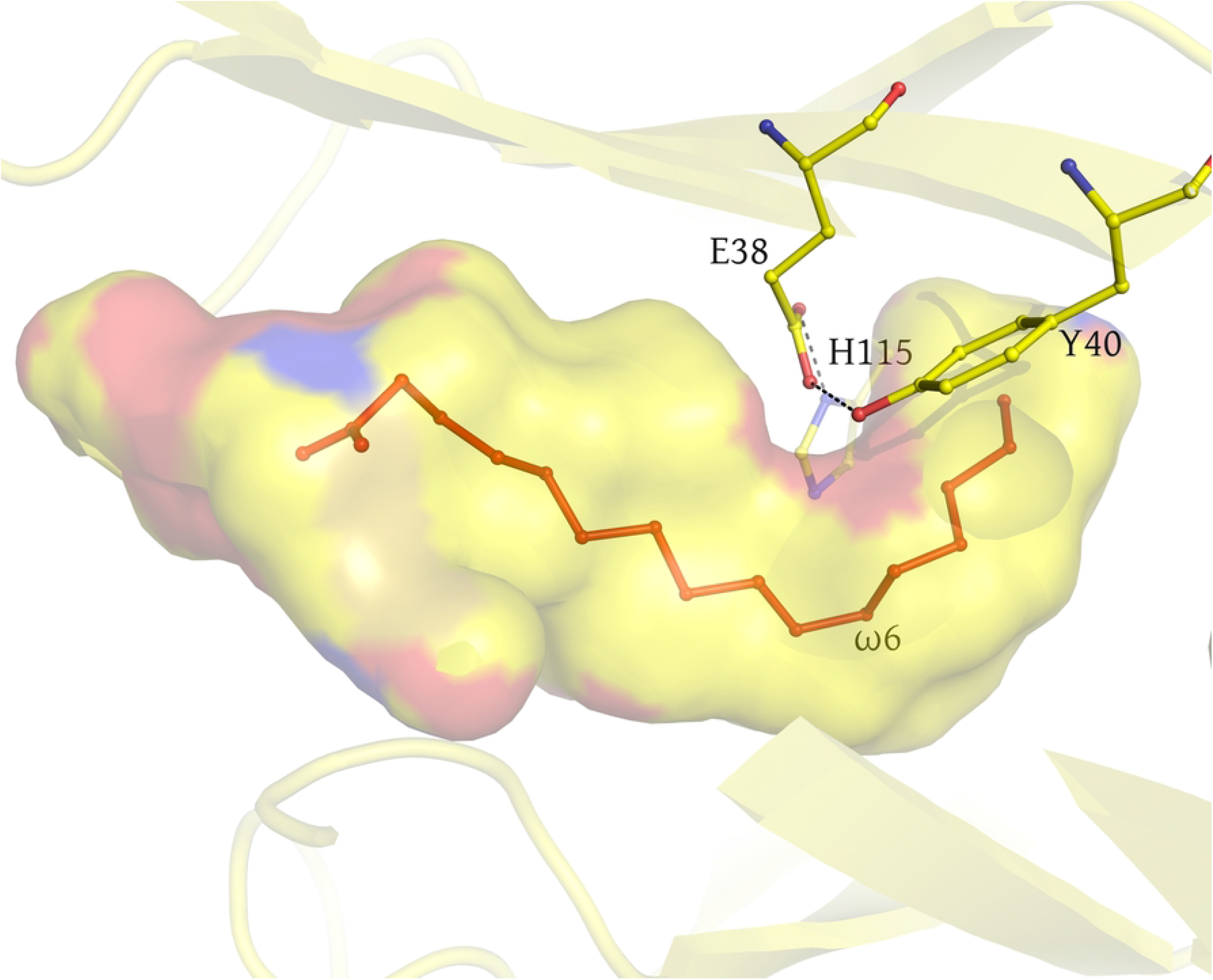
The kink in Lili-Mip2 binding pocket. E38, held in place by Y40 and H115, shapes the bend near the ω6 of the fatty acid(red). The figure was made using the deposited structure with PDB-ID 4NYQ.

Previous work in our lab has shown that Lili-Mip2 is a thermophilic protein that does not denature even at 100 °C [10]. Another lipocalin, β-lactoglobulin, is thermodynamically stable at low pH. There is a difference of 16.6 °C in melting temperature (T_m_) of β-lactoglobulin as the pH changes from 7.5 to 1.0 [25]. The stability of lipocalins at low pH could be attributed to glycosylation. This study aims to elucidate the stability and binding properties of recombinantly expressed Lili-Mip1, 2, and 3 and compare these factors to the properties of the Lili-Mips purified from the cockroach midgut. The results provide detailed information to engineer ligand binding by mutating residues in the active site.

## 2 MATERIALS AND METHODS

### 2.1 Cloning

Lili-Mip1, Lili-Mip2, and Lili-Mip3 were synthesized by GeneArt and cloned in pYES2-CT vectors with a C-terminal Histidine tag [8,10]. Further, Lili-Mip1, 2, and 3 were sub-cloned into the pYES2 vector without an affinity tag. All constructs were codon optimized for yeast expression and had an N-terminal secretion signal (Ost1 or α-factor) for protein secretion into the media [26]. A point mutation in Lili-Mip2, at residue 38, from glutamate to alanine at 38 (Lili-Mip2-E38A) was made by site-directed mutagenesis. Lili-Mip2 in the pYES2 vector was modified using forward primer (5’ GTTTCTAACATCACC gct TTCTACTCTGCTCATG 3’) and reverse primer (5’ GAGCAGAGTAGAA AGC GGTGATGTTAGAAAC 3’) through PCR and the modified vector was transformed into yeast and sequence verified by Sanger sequencing.

### 2.2 Expression and purification

*Saccharomyces cerevisiae* strain of FGY217 (*MATα*, ura3-52, *lys2_201*, and *pep4*) was used to express and secrete the protein. Yeast was transformed with Lili-Mip plasmid by the Lithium acetate method [27]. The transformed yeast colonies were inoculated in media lacking uracil with 2% Dextrose and grown overnight at 30 °C, 220 rpm. The Optical Density (OD) value at 600 nm was measured, and the culture was diluted with new YPD media to an OD of 0.2/ml. This was again grown till the OD reached 0.6-0.7, induced with 2% galactose, and incubated for 24 hours at 30 °C, 220 rpm. The supernatant was clarified by centrifugation at 4000 ×g for 10 minutes. The clarified media was concentrated using Sartorius Vivaflow200, with a 10kDa molecular weight cut off filter to a final volume of 500ml. The media was buffer exchanged into 50mM sodium acetate at pH 5.0 and 10mM NaCl while concentrating the media. The buffer exchanged media was purified using cation exchange chromatography on the SP FF column (Cytiva). The Lili-Mip protein eluted between 100mM to 200mM of NaCl (on a gradient that extended to 1.0M NaCl). The protein fractions were concentrated using a 10 kDa filter in an Amicon spin filtration system. The concentrated protein was purified using the size exclusion chromatography column S75 (Prep Grade - Cytiva) in 50mM Sodium acetate pH 5.0 and 50mM NaCl buffer. The Lili-Mip proteins resolved as single peaks after size exclusion chromatography.

### 2.3 Lipid Extraction and Delipidation of Lili-Mip

Lili-Mip was delipidated using the Bligh Dyer method [29]. The chloroform phase containing the fatty acids was dried and used for fatty acid analysis using LC-MS/MS. The aqueous phase contained the delipidated protein, and it was dried and resuspended in an appropriate buffer for binding studies.

### 2.4 Intrinsic Tryptophan fluorescence

Fluorescence measurements were carried out in 0.05 M Na-acetate buffer pH 4.7 or 0.05 M Na-phosphate buffer pH 4.7/7.2/8.2, both containing 100mM NaCl buffer. Delipidated Lili-Mip1,2 or 3 were diluted to 4 μM in the buffer, and ligand (dissolved in ethanol) was added in the concentration range from 0-35 μM. The protein solutions were incubated for 1 min at room temperature (25 °C). The samples were excited at 290 nm (slit 2.5 nm), and the emission was measured at 325 nm (slit 2.5 nm) on a Cary fluorescence spectrophotometer. The analysis of the data was performed as described earlier [28].

### 2.5 Targeted analysis of fatty acids using LC/MS/MS

An Agilent 1290 Rapid Resolution liquid chromatography (LC) system coupled to an Agilent 6470 series QQQ mass spectrometer (MS/MS) was used to analyze fatty acids in each sample (Agilent Technologies, Santa Clara, CA). The methods are similar to what is described in the paper by Yang, Adamec, and Regnier [30]. The internal standard for the assay was the D_3_-CMP derivatized pure fatty acid. Each sample was spiked with 50 ng of the internal standard before analysis. An Agilent Eclipse Plus C-18 (2.1 mm x 50 mm) 1.8 µm column was used for LC separation. The buffers were (A) water + 0.1 % formic acid and (B) isopropanol/acetonitrile (50/50 v/v) + 0.1% formic acid. The linear LC gradient between 10% and 100% of B was used. The post time was set to 6 minutes. The flow rate was 0.3 mL/min. Multiple reaction monitoring was used for MS analysis. The data were acquired in positive electrospray ionization (ESI) mode. The jet stream ESI interface had a gas temperature of 325°C, a gas flow rate of 8 L/minute, a nebulizer pressure of 45 psi, a sheath gas temperature of 250°C, a sheath gas flow rate of 7 L/minute, a capillary voltage of 4000 V in positive mode, and nozzle voltage of 1000 V. The ΔEMV voltage was 300 V. Agilent Mass hunter Quantitative analysis software was used for data analysis (version 10.1).

Native crystal isolation from the gut crystals was carried out, as described previously by Banerjee et al. [8]. Native Lili-Mip crystals were extracted in 50% PEG400, and the samples were shipped in 50% PEG400. Further, the crystals were dissolved in appropriate buffers for biochemical studies.

### 2.6 pH Probing in the gut

pH levels were assessed using a Thermo Scientific Orion micro-pH probe based on a protocol adapted for cockroaches from tsetse flies [31]. Specifically, females were dissected to remove embryos during the late stages of lactation. The embryos were dissected to remove the gut. The guts were ruptured and centrifuged through a fine mesh (polyester, 150 mesh) filter at 1,000 RPM for two minutes to separate the gut contents from gut. The pH of the contents of the gut lumen (containing Lili-Mips), the cells of the gut (homogenized), and the remaining body parts (homogenized) were determined.

### 2.7 Docking Calculations

CANDOCK (version 0.6.0), an in-house docking software, was used to generate docking conformations of all fatty acids with Lili-Mip2 (PDB ID: 7Q02), including a selection of the binding site. The fatty acid structures for myristic acid (14:0), palmitoleic acid (16:1), and linoleic acid (18:2) were drawn on BioChemDraw, cleaned in 3D, and saved as a Mol2 file for docking. CANDOCK was used with default parameters, and the radial-mean-reduced-6 (RMR6) was used as the “Selector” parameter for docking to select the top pose as benchmarked previously [32]. The docking scores of all 96 different potential energy functions were calculated for all docking states. Protonation states of the protein bound to the fatty acids were determined for four pH values: 4.7, 5.9, 7.2, and 9.1. The distance from the last carbon atom on the fatty acid to carbon CH_2_ on W20 in the binding pocket was calculated using VMD software for each docked pose; the fatty acid carbon atoms, C14 on myristic acid, C16 on palmitoleic acid, C18 on linoleic acid were used [33].

### 2.8 Molecular Dynamics Simulations

Docked pose of the fatty acid with the lowest distance (3-5Å) from carbon CH2 on W20 was used for Molecular Dynamics (MD) simulations with GROMACS [34]. MD calculations were performed for 200 ns with the docked conformation of each fatty acid with protein surrounded in explicit water molecules. We used CHARMM-26 force field and TIP3P water models in a 42 Å cubic box with periodic boundary conditions [35,36]. To prepare the system for MD, we ran 50,000 steps of steepest descent energy minimization with an initial step size of 0.1 Å. The particle mesh Ewald (PME) scheme was used for long-range electrostatics [37] using the Coulomb cutoff distance of 1.2 Å. A van der Waals (VdW) cutoff type was specified with a cut-off distance of 1.2 Å. All other parameters in energy minimization were set to the default values in GROMACS. After energy minimization, a 100 ps MD simulation was run in the NVT ensemble (constant number of molecules, volume, and temperature) using the Berendsen thermostat at a reference temperature of 300 K, sampled at every 0.1 ps with a chain length of 4 for equilibration. The electrostatic cutoff distance was set to 1.5 nm. To get the isothermal-isobaric NPT ensemble (constant number of molecules, pressure, and temperature), we used a time step of 0.002 ps with Berendsen thermostat at a reference temperature of 298 K and a time constant of 1 ps for the entire system and the Berendsen barostat [38] at a reference pressure of 1 bar with a time constant of 2 ps and a compressibility value of 4.5e-5 bar^-1^. A final production MD simulation was done for 200 ns with initial velocities generated by a Maxwell distribution at 298 K. The fatty acid was not constrained during the equilibration and production MD run. All other parameters in the MD run were set to default values used by GROMACS.

### 2.9 Deglycosylation using PNGase

The recombinant Lili-Mips were treated with PNGase F (New England Biolabs) to remove the N-linked oligosaccharides. Glycosylated Lili-Mip and PNGase F were mixed in a buffer having 1X GlycoBuffer-2 (0.5 M Sodium phosphate pH 7.0, 0.2 M EDTA). The reaction mixture was incubated at 37°C for 24 hours. SDS-PAGE was used to differentiate glycosylated and deglycosylated Lili-Mip.

### 2.10 Crystallization

The Lili-Mips were concentrated using 10kDa Amicon filters. The calculated extinction coefficient of 28100 M^-1^cm^-1^ was used to measure the concentration using a NanoDrop 2000 (Thermo Fisher) spectrometer. Hanging drops were set up using a Mosquito robot (200nl protein + 200nl well solution with a 50ul well solution). Lili-Mip2-E38A crystals were obtained in 0.002 M Zinc sulfate heptahydrate, 0.08 M HEPES pH 7.0; 25 % v/v Jeffamine® ED-2003 as the precipitant at 20 °C. Lili-Mip1 crystals were obtained in 0.2M Potassium Nitrate pH 6.9; the precipitant was 20%(W/V) PEG 3350. The crystals were cryo-protected by adding 5% glycerol.

### 2.11 Data Collection, Structure solution, and Refinement

The data for the Lili-Mip2-E38A mutant was collected at the MBC-CAT beamline at the Advanced Light Source at Berkeley (900 frames at 0.2-degree oscillation). The data were processed using XDS [39] and scaled using the Aimless package in the CCP4-suite [40,41]. The data were good to 2.95 Å resolution. The structure was determined by molecular replacement with the deposited structure of Lili-Mip (PDB ID 7BKX). The maps clearly showed the absence of the side chain at position 38. The structure was refined using Refmac to a final R-factor and R-free of 23.4 and 29.4, respectively [42]. Statistics of data collection, refinement, and structure quality are in Supplementary Table 1. The data for Lili-Mip1 was collected at the APS GMCA-CAT beamline ID 23D-D. Data processing with XDS and scaling suggested good data to 2.16 Å resolution. The structure determination was by molecular replacement using the deposited 4NYQ as the model. The refined model had a final R-factor and R-free of 0.19 and 0.25, respectively. For both structures, iterative model building was carried out with the program Coot [43].

**Table 1.** Binding affinity of Lili-Mip1, 2 and 3 with different fatty acids. Intrinsic Tryptophan fluorescence is employed to identify binding affinities of Lili-Mip1, 2 and 3 for different fatty acids. All values are in μM. The protein is in 25mM Sodium acetate buffer pH 4.8, 100mM NaCl. The experiments are repeated with three biological replicates, n=3 and are drawn as mean ± S.D.

| In $\mu\text{M}$ | Myristic acid (14:0) | Myristoleic acid (14:1) | Palmitic acid (16:0) | Palmitoleic acid (16:1) | Stearic acid (18:0) | Oleic acid (18:1) | Linoleic acid (18:2) |
| --- | --- | --- | --- | --- | --- | --- | --- |
| Lili-Mip1 | 1.96 ±0.83 | 2.06 ±0.42 | 1.76 ±0.88 | 1.54 ±0.91 | 1.71 ±0.56 | 0.93 ±0.31 | 1.56 ±0.33 |
| Lili-Mip2 | 2.44 ±2.68 | 2.78 ±0.50 | 1.41 ±0.88 | 2.30 ±1.32 | 2.32 ±1.32 | 1.71 ±1.19 | 2.19 ±1.41 |
| Lili-Mip3 | 1.60 ±0.51 | 0.68 ±0.49 | 0.82 ±0.61 | 0.88 ±0.62 | 1.78 ±1.74 | 0.24 ±0.11 | 0.27 ±0.08 |

### 2.12 Thermal stability Assay

Tycho NT.6 (Nano Temper) was used to measure the melting temperature (T_m_) of recombinant Lili-Mips and its mutants. The protein stability was measured in different buffers (50mM HEPES 7.4 with salt ranging from 0.5M to 2M, 50mM Sodium phosphate buffer ranging from pH 4.7 to 9.3) to determine the pH and salt dependence on stability. The protein concentration was set to 0.1mg ml^-1^. The heating range was between 35 °C and 95 °C. Model figures and plots were made with Pymol [44] and R [45], respectively.

## Results and Discussion

Lili-Mip1, 2 and 3 are thermophilic proteins at low pH (4.8). Lili-Mip2 is the most thermostable, followed by Lili-Mip1 and 3. Upon increase in the pH, the stability of Lili-Mip1,2 and 3 decreased markedly. Lili-Mip2 had the maximum melting temperature(ΔT_m_) change of around 50 °C on pH change from 4.8 to 9.3 (Figure 2A). The thermostability was independent of the amount of salt in the environment (0.5M to 2M NaCl), indicating that the Lili-Mip1, 2, and 3 are also stable at high salt (Figure 2B). Native Lili-Mip also shows similar behavior. Though Lipocalins are known to be thermostable, such a large change in T_m_ has not been observed so far. Lacto-globulin, a bovine milk protein, shows a change in thermal stability (ΔT_m_ =16 °C) under similar conditions [25]. Other Lipocalins were shown to have no such pH-dependent behavior (Supplementary Figure S1). In order to understand where the stability stems from, we deglycosylated Lili-Mip1. Deglycosylated Lili-Mip1 showed a marginal decrease in stability (Supplementary Figure S2). However, the removal of glycans did not affect the pH-dependent change in the thermal stability of Lili-Mip1. The effect of delipidation of Lili-Mip1, 2, and 3 were investigated. The delipidated Lili-Mip1 and 3 did not show any difference from the holo-protein. However, there is a 10°C decrease in the stability of Lili-Mip2 at low pH (Supplementary Figure S3). We observed Lili-Mip2-E38A with a mutation of glutamate to alanine in the binding pocket is more thermostable even at basic pH (Supplementary Figure S4). We concluded that ligand binding to Lili-Mip is probably not linked with the pH-dependent effect on stability.

**Figure 2.**
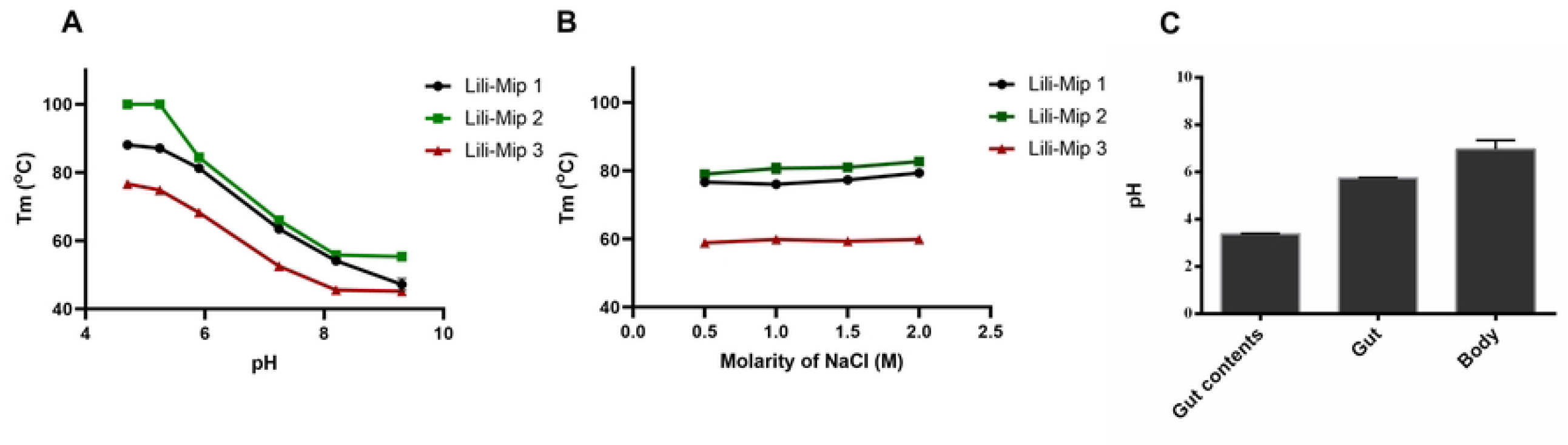
Temperature dependent denaturation of recombinant Lili-Mip 1,2 and 3 with respect to pH and salt. (A) Left -Melting temperature (T_m_) of Lili-Mip1,2 and 3 in 50mM sodium phosphate buffer at pH 4.7, 5.2, 5.9, 7.2, 8.2, 9.3 as mentioned in methods. (B) Melting temperature (T_m_) of Lili-Mip1, 2 and 3 in 50mM HEPES pH 7.4 at different salt concentrations from 0.5M, 1M, 1.5M and 2M. Data was collected on TychoNT.6 (Nanotemper). The experiments are repeated with three biological replicates, n=3 and are represented as mean ± S.D. (C) pH of the gut contents (primarily Cockroach milk), cells of the gut and body measured using a microprobe shows Lili-Mip is stored in acidic conditions whereas the cells are at a higher pH. The experiments are repeated with two cockroaches(n=2) and are represented as mean ± S.D.

To understand the physiological relevance of this, we measured the pH of the embryo midgut and the cells of the embryo. We show that the gut contents, which are essentially cockroach’s milk containing Lili-Mips have an acidic pH of 3.38. However, the gut cells showed a pH of 5.76. The remaining body showed a pH of 6.96 (Figure 2C). The cells are maintained in a basic environment as compared to the milk. This provides one possible explanation for this behavior. The embryo ingests Lili-Mip into its midgut, where they are stored until required. The increased stability of Lili-Mips at an acidic pH enables the long-term storage of the protein and its ligand (fatty acid). When the cells of the embryo need food, the Lili-Mip can be taken into the cells and assimilated. The decreased stability in higher pH allows it to be broken down and used as food. The Lili-Mips shown in the stability assay are all recombinantly expressed and purified from Baker’s yeast but are likely to represent the stability of naturally occurring Lili-Mip structures.

To find what lipids are bound to native or recombinant Lili-Mips, we used a fatty acid extraction coupled with Mass Spectrometry. This showed Lili-Mip1 binds primarily to palmitoleic acid (16:1), palmitic acid (16:0), oleic acid (18:1), and stearic acid (18:0). Lili-Mip2 binds to palmitic acid (16:0), palmitoleic acid (16:1), stearic acid (18:0) and oleic acid (18:1). Lili-Mip3 binds to palmitic acid (16:0), stearic acid (18:0), palmitoleic acid (16:1), oleic acid (18:1). Native Lili-Mip had palmitic acid (16:0), stearic acid (18:0), palmitoleic acid (16:1) and linoleic acid (18:2) as the dominant fatty acids (Figure 3). These data suggest that recombinant Lili-Mip from yeast binds to fatty acids with 16-18 carbon atoms. Contrary to our initial hypothesis, there does not seem to be a preference for the degree of saturation. It is important to note that 16 and 18-carbon fatty acids are the predominant fatty acids in yeast. Native Lili-Mip also follows the trend, the only difference being the increased presence of linoleic acid. This could be attributed to the fact that Linoleic acid is known to be present in *Diploptera punctata* but not in *Saccharomyces cerevisiae* [8]. These results suggest that the abundance of fatty acids bound to Lili-Mip depends on the endogenous composition of the fatty acids in its milieu. In other words, Lili-Mip1, 2, and 3 show similar specificity to fatty acids (Figure 3). The composition of different Lili-Mips differs during the gestation of the cockroach. This could be one way the mother cockroach can regulate the supply of fatty acids to the embryo during development. The fact that Lili-Mip1, 2, and 3 could transport a wide range of fatty acids shows that the embryo can have a constant supply of fatty acids as nutrition.

**Figure 3.**
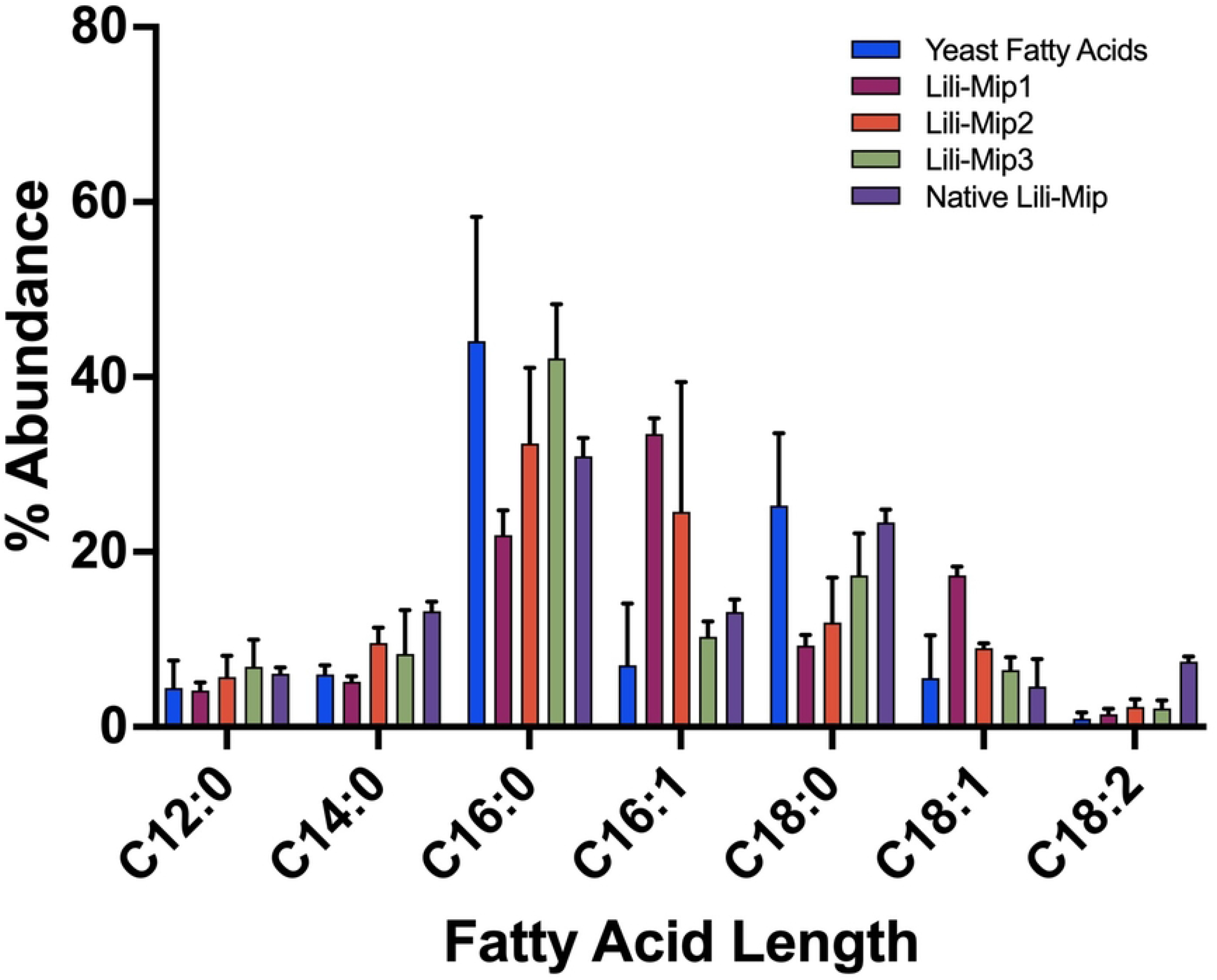
Fatty acid profile of recombinant and native Lili-Mips using LC/MS/MS. (A) Fatty acids bound to Lili-Mip1, 2, 3 and native Lili-Mip crystals. (B) Fatty acid profile of Lili-Mip2 and Lili-Mip2-E38A. Each fatty acid peak density was compared with the corresponding standard. The experiments are repeated with three biological replicates, n=3 and are drawn as mean ± S.D.

We show that native or recombinant Lili-Mips can bind to different fatty acids. We exploited the single tryptophan at the base of the ligand binding site to measure how tightly these fatty acids bind to Lili-Mip1, 2, and 3. The fluorescence of this tryptophan gets quenched on ligand binding. Intrinsic tryptophan fluorescence was employed to find the binding affinity of different fatty acids to Lili-Mip1, 2, and 3 (Supplementary Figure S5). Fatty acids of varying chain lengths (14, 16, 18) and degrees of unsaturation (0, 1, 2) were assayed. All tested fatty acids bind with micromolar range affinities to the different Lili-Mips (Table 1). This data suggests that the length of the acyl chain and the unsaturation does seem to significantly affect the affinity of binding. This reiterates that the fatty acids the embryo receives almost exclusively depend on the fatty acid composition of the milk-secreting cells.

We considered how an increase in pH could affect the binding affinity. Binding studies with Lili-Mip1 against palmitic acid showed that Lili-Mip binds to the ligand with similar binding affinities between pH 4.7 and 8.2. This also reaffirms that a change in thermal stability with pH is most likely not a consequence of ligand binding/unbinding. Lili-Mip2-E38A did not show any quenching of fluorescence on binding to the ligand. Therefore, we could not measure the binding affinity of ligand binding even though fatty acid analysis showed ligand binding. From the structure of the protein with the E38A mutation, the change in orientation of the Phenyl alanine’s (98 and 100), makes the Tryptophan inaccessible to the fatty acid chain of the substrate. This would suggest that the environment of Trp-20 does not change on fatty acid binding.

The structure of Lili-Mip1(PDB ID: 8F0Y) and Lili-Mip2-E38A (PDB ID: 8F0V) showed a density for the bound fatty acid. However, it was difficult to ascertain the length and nature of fatty acid bound. The biochemical data indicated that these proteins could bind to multiple fatty acids. This correlates with the lack of clear density in the crystallographic data suggesting heterogeneity in fatty acid binding among the molecules that make up the lattice. This is not surprising as the residues that make crystal contacts are far away from the residues in the binding pocket.

In the Lili-Mip1 structure, we found that Phe-98 and Phe-100 have moved upwards, causing a decrease in the size of the ligand binding pocket (Figure 4A-D). To better understand how the ligand binding pocket size changes between structures, we quantitated the cavity volumes of different Lili-Mip using parKVfinder [46]. This is seen better in contrast with Lili-Mip2, where the Phe-98 and Phe-100 face downwards, increasing the cavity volume (Figure 4E-G). The decrease in the size of the binding pocket seems directly correlated to the side chain orientation of Phe-98 and Phe-100.

**Fig 4.**
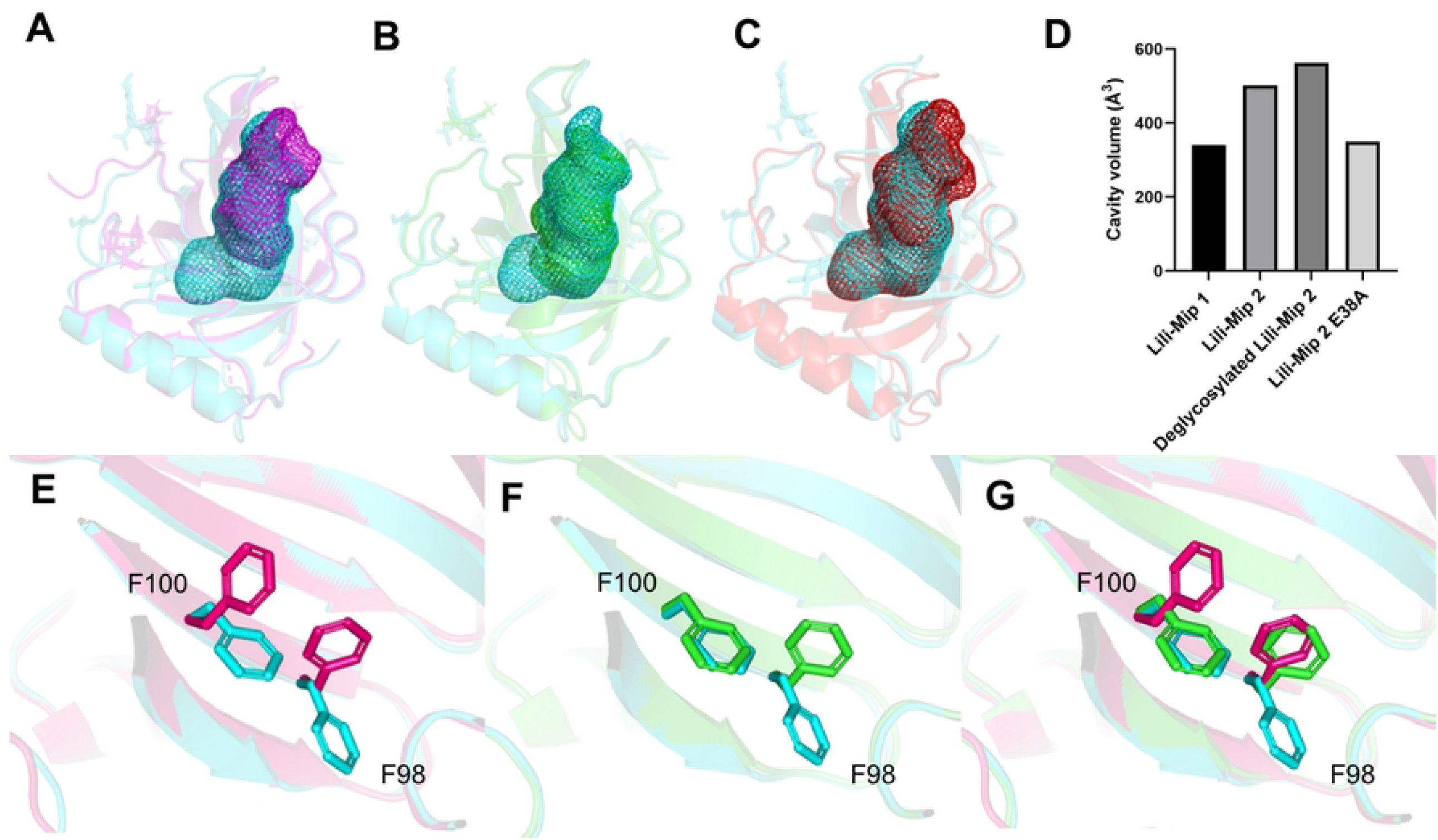
Cavity volumes of Lili-Mip1 (8F0Y), Lili-Mip2 (7BKX) Deglycosylated Lili-Mip2 (7Q02) and Lili-Mip2-E38A (8F0V) Parkvfinder was used to find cavity volumes(mesh) in Lili-Mips in order to see the effect of F98 and F100 on ligand binding (A) Lili-Mip2 (cyan) superimposed to Lili-Mip1 (magenta) (B) Lili-Mip2 (cyan) superimposed to Lili-Mip2-E38A (C) Lili-Mip2 (cyan) superimposed to Deglycosylated Lili-Mip2 (red) (D) Histogram showing the different Lili-Mips and its mutant and its cavity volumes F98 and F100 assumes different conformations in the crystal structure. The confirmations are shown as an overlay of (E) Lili-Mip2 (cyan) against Lili-Mip1 (magenta) (F) Lili-Mip2 against Lili-Mip2-E38A (green) (G) Superposition of Lili-Mip2, Lili-Mip1 and Lili-Mip2-E38A.

In Lili-Mip2-E38A, a zinc atom is present as an artifact of crystallization. Zinc interacts with the carboxylic part of the ligand and with residues D53, E61, and H63, like what happens with glycosylated Lili-Mip2 (PDB ID:7BKX) [10]. The structure of Lili-Mip2 showed that in the ligand binding pocket, E38 is held in position by hydrogen bonds from H115 and Y40, enabling ligand binding [10]. The mutation of this glutamate to alanine was predicted to influence ligand binding. On superimposition with wild-type Lili-Mip2 structure, the cavity available for ligand binding decreases (Figure 4B). The F98 position has moved upwards, causing this upward shift, whereas F100 is unaltered (Figure 4F). The mutation to alanine only slightly altered the position of Y40 and H115.

Deglycosylation of Lili-Mip2 does not change the position of F98 and F100, and the cavity volumes are not affected as much (Figure 4C). This indicated that glycosylation might not have many roles in regulating the size of the hydrophobic cavity. We conclude that the position of F98 and F100 has a bearing on the cavity volumes. The volume of the binding pocket decreases in Lili-Mip2-E38A as opposed to Lili-Mip2 (Figures 4B and 4D). The cavity volume does not change much when Lili-Mip2 is deglycosylated (Figures 4C and 4D). The orientation of these residues towards the entrance of the binding pocket in Lili-Mip1 shows that similar composition of ligand binding pocket causing a dramatic change (221.75 Å³) compared to Lili-Mip2 (Figures 4A and 4D). The three unique conformations of F98 and F100 in Lili-Mip1, Lili-Mip2, and Lili-Mip2-E38A correspond to different cavity volumes. The different conformations of F98 and F100 allow Lili-Mips to bind fatty acids of different lengths, thereby explaining promiscuity.

MD simulations were performed to understand the ligand binding properties of deglycosylated Lili-Mip2 at different pH from 4.7 to 9.1. The distance between the invariant W20 and the last Carbon of fatty acid was calculated to find the fatty acid’s location over the simulation’s time period. The probability density plot for fatty acid binding in deglycosylated Lili-Mip2 shows that at a pH of 4.7 the fatty acid stays in the ligand binding pocket for a longer time than it does at the pH inside the cells (Figure 5). Interestingly as the pH increases to 9.1, probability density plots begin to suggest that binding is tighter. This result, along with the lower stability of the protein at physiological pH supports the idea that once absorbed into the cells, the protein can be broken down, and the fatty acid, amino acids, and sugars can be used. The simulations show that myristic acid binds farther away from W20 than palmitoleic acid. This is expected due to the shorter length of the acyl chain. Similar to that observed for other lipocalins, the simulations also suggest that the residues that change conformation the most are at the entrance of the binding pocket (Supplementary Figure S6)

**Figure 5.**
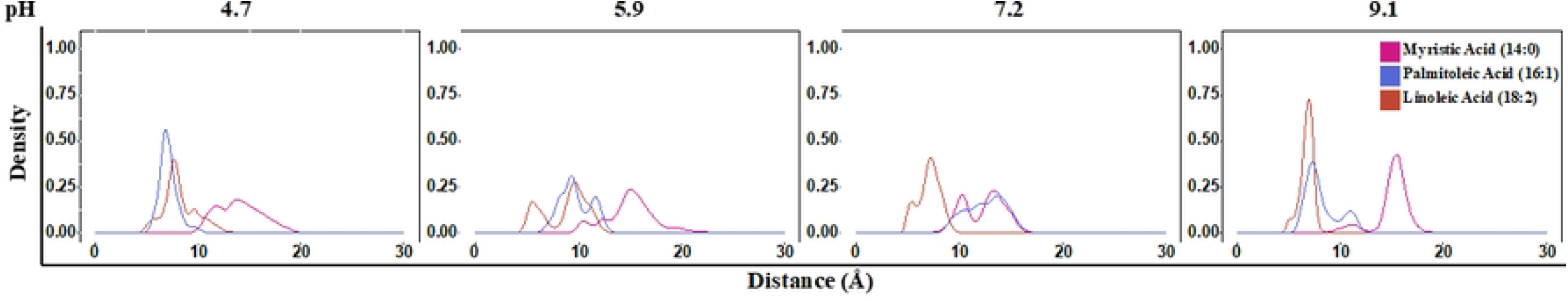
Molecular Simulations of deglycosylated Lili-Mip 2(PDB ID: 7Q02) shows nature of fatty acid binding. Probability density distance plots myristic acid (pink), palmitoleic acid (blue) and linoleic acid (orange) at different pH.

In summary, results from our studies on fatty acid binding affinity, promiscuity, structures of Lili-Mip1 and Molecular dynamics simulations all show that Lili-Mips bind with similar affinity to several fatty acids. This allows them to carry different fatty acids, and the population of the fatty acid delivered as food can be controlled by the amounts of the different fatty acids in in the cells that produce the protein. Lili-Mip is a highly thermostable protein, and the stability is pH dependent. We hypothesized that this allows the protein-lipid-glycan complex to be very stable in the gut and less stable once inside the cell where the pH is neutral. Measurements of pH in the gut and cells confirm the difference in pH in the cockroach. This would allow the cells to break down Lili-Mips and use them as a source of metabolites - amino acids, fatty acids, and sugars – to promote embryo growth within the brood sac. The different orientations of Phe-98 and Phe-100 control the binding pocket volume and allow the binding of different chain-length fatty acids to bind with similar affinities.

## Acknowledgements

We thank ALS(ALS 4.2.2 beamline) and APS(APS 23-ID-D beamline) synchrotrons for providing the beamtime for data collection. We would like to thank Jay Nix, Beamline Director for the Molecular Biology Consortium, for his help with data collection. We thank the Chemical Genomics Facility at Purdue University for using Tycho. All Mass-spectrometry data were collected at the Bindley Bioscience Center, Purdue University.

## Data Availability

The protein structures are deposited in the protein data bank with PDB ID 8F0V and 8F0Y.

The authors declare no competing interests.

